# Conformational specificity of opioid receptors is determined by subcellular location irrespective of agonist

**DOI:** 10.1101/2021.03.18.435866

**Authors:** Stephanie E. Crilly, Wooree Ko, Zara Y. Weinberg, Manojkumar A. Puthenveedu

## Abstract

The prevailing model for the variety in drug responses is that they stabilize distinct active states of their G protein-coupled receptor (GPCR) targets, allowing coupling to different effectors. However, whether the same ligand can produce different GPCR active states based on the environment of receptors in cells is a fundamental unanswered question. Here we address this question using live cell imaging of conformational biosensors that read out distinct active conformations of the δ-opioid receptor (DOR), a physiologically relevant GPCR localized to Golgi and the surface in neurons. We show that, although Golgi and surface pools of DOR regulated cAMP, the two pools engaged distinct conformational biosensors in response to the same ligand. Further, DOR recruited arrestin on the plasma membrane but not the Golgi. Our results suggest that the same agonist drives different conformations of a GPCR at different locations, allowing receptor coupling to distinct effectors at different locations.

## Introduction

A given GPCR can generate a diverse array of signaling responses, underscoring the physiological and clinical relevance of this class of proteins. Endogenous and synthetic ligands which “bias” the responses of a given receptor towards one response or another are a key aspect of this signaling diversity (*1*). This diversity in responses provides several opportunities to target specific GPCR signaling responses to reduce potential adverse effects while managing a variety of clinical conditions. However, using bias to precisely tune GPCR signaling has been difficult, suggesting that we are missing some key piece in our understanding. Understanding the cellular mechanisms that contribute to individual components of the integrated signaling response is therefore of profound importance to understanding GPCR pharmacology.

The specific conformations adopted by GPCRs, which preferentially allow coupling to distinct effectors, is likely a key determinant of which specific downstream signaling response is amplified (*2*–*5*). Recent studies support an allosteric model of coupling in which binding to both agonist and G protein stabilizes an active state of the receptor, among a number of states which a given receptor may sample (*5*–*9*). In addition to canonical G protein effectors, β-arrestins interact with GPCRs through additional receptor conformations and can serve as scaffolds for kinase signaling pathways (*10*–*12*). However, efforts to develop compounds which stabilize one set of conformations and therefore bias the receptor response to specific pathways have been promising but difficult to translate to *in vivo* models (*13*–*15*).

One hypothesis that could explain this difficulty is that the same agonist could drive coupling of the same receptor to different core signaling proteins based on the immediate subcellular environment of receptors. While an exciting idea with profound implications, this hypothesis has been difficult to test using traditional methods, because receptor signaling readouts are complex and have been difficult to separate based on location.

Here we use the δ-opioid receptor (DOR), a physiologically and clinically relevant GPCR, as a model to test this hypothesis. DOR localizes primarily to the Golgi in neuronal cells, with a small amount on the plasma membrane (*16*–*22*). DOR can be activated both on the plasma membrane and the Golgi by synthetic agonists, but whether the two activation states are different is not clear (*23*). Relocating DOR from intracellular compartments to the plasma membrane (PM) increases the ability of DOR agonists to relieve pain (*17, 21, 24, 25*), illustrating the importance of understanding whether DOR activation on the Golgi is different from that on the plasma membrane.

Here we leverage conformational biosensors and high-resolution imaging to test whether DOR activation on the Golgi is different from that on the plasma membrane. We show that DOR on the PM, when activated by the selective DOR agonist SNC80, can recruit both a nanobody-based sensor and a G protein-based sensor that reads out active DOR conformations, as well as β-arrestins. In contrast, DOR in the Golgi apparatus, when activated by the same ligand, recruits the nanobody sensor, but not the G protein-based sensor or β-arrestins. Nevertheless, Golgi-localized DOR is competent to inhibit cAMP. Together these data demonstrate that these biosensors could be used to read out subtle differences in GPCR conformations even if signaling readouts are similar. Our results that the downstream effectors recruited by the same GPCR, activated by the same ligand, depend on the location of the receptor, suggest that subcellular location could be a master regulator of GPCR coupling to specific effectors and signaling for any given GPCR-ligand pair.

## Results

Nanobody and miniG protein biosensors are emerging as powerful tools to study the effects of ligand-induced receptor conformational changes at the molecular and cellular level (*26*–*28*). These sensors differentially engage the μ-opioid receptor (MOR) and κ-opioid receptor (KOR) activated by different agonists *in vitro* (*29*) or in the PM (*30*), suggesting that they can provide a readout of conformational heterogeneity in agonist-stabilized active states. Because DOR is highly similar to MOR and KOR in the intracellular regions recognized by the sensors (*31*), we asked whether these sensors could be optimized to read out specific active DOR conformations at distinct subcellular locations. We used the nanobody biosensor, Nb39 which recognizes opioid receptor active conformations through residues conserved across MOR, KOR, and DOR (*32, 33*), and the miniGsi biosensor, which mimics the interaction of the Gαi protein with GPCRs (*28, 34*), as two orthogonal readouts of DOR conformations.

As an initial step, we first tested whether these sensors report active DOR conformations on the PM, similar to what has been reported for MOR and KOR. We used total internal fluorescence reflection microscopy (TIR-FM), which uses an evanescent wave to specifically excite fluorescent proteins on the PM to a depth of approximately 100nm into the cell (*35*), to visualize sensor recruitment to activated DOR on the PM with high sensitivity (Fig. 1A). When cells were treated with either small molecule agonist SNC80 or peptide DPDPE, Nb39 and miniGsi were rapidly recruited to DOR on the plasma membrane (PM DOR), as observed by a rapid increase in fluorescence (Fig. 1B-E). Fluorescence of both sensors increased significantly after treatment with either agonist, but not inverse agonist ICI174864 (ICI) (Fig. 1D-F), indicating specificity of both sensors for an agonist-induced active conformation. Interestingly, DPDPE recruited Nb39 to PM DOR more strongly than SNC80 (Fig. 1D,F), whereas the opposite trend was observed for miniGsi (Fig. 1E,F), suggesting that these sensors might report selective conformations of DOR induced by different agonists. MiniGs, which mimics Gαs protein interaction with Gs-coupled GPCRs, was not recruited to PM DOR (Fig. 1F), suggesting that the miniGsi sensor specifically reports activation of the Gi-coupled DOR. DOR expression levels were overall comparable across conditions (Fig. S1A). Overall, our data show that Nb39 and miniGsi report agonist-induced active conformations of PM DOR.

**Fig. 1.**
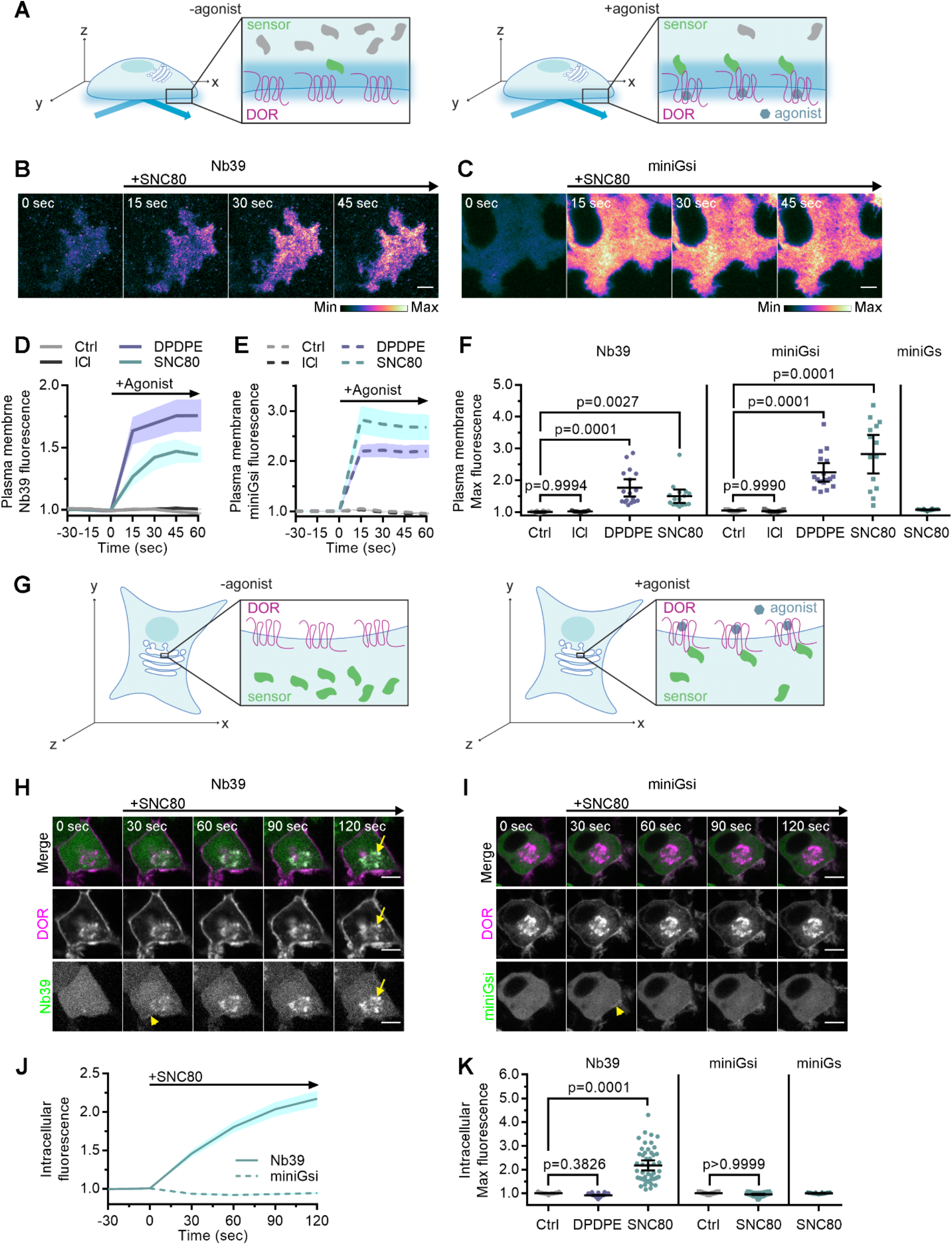
Nb39 and miniGsi are differentially recruited to plasma membrane and intracellular DOR. **(A)**. Schematic of biosensor recruitment to DOR in the PM using total internal reflection fluorescence microscopy (TIR-FM). Only fluorescent proteins within the evanescent wave close to the PM are excited, such that baseline fluorescence is low when biosensors are diffuse in the cell but increases upon agonist addition as biosensors recruited to active DOR in the plasma membrane. **(B)**. Nb39-mVenus in PC12 cells expressing SNAP-DOR imaged using TIR-FM to capture recruitment to the PM after addition of 10µM SNC80 (scale bar = 5µm). **(C)**. Venus-miniGsi in PC12 cells expressing SNAP-DOR imaged using TIR-FM to capture recruitment to the PM after addition of 10µM SNC80 (scale bar = 5µm). Calibration bars indicate relative fluorescence values in scaled images. **(D)**. Increase in Nb39-mVenus fluorescence by TIR-FM normalized to the mean baseline fluorescence over time after addition of 10µM DOR agonist, DPDPE or SNC80, 10µM inverse agonist, ICI174864 (ICI), or vehicle control (Ctrl, n=10 cells; ICI, n=17 cells; DPDPE, n=17 cells; SNC80, n=16 cells; all across 3 biological replicates defined as coverslips prepared and imaging independently; solid line indicates mean, shading +/− SEM). **(E)**. Increase in Venus-miniGsi fluorescence by TIR-FM normalized to the mean baseline fluorescence over time after addition of 10µM DPDPE or SNC80, 10µM inverse agonist ICI, or vehicle control (Ctrl, n=17 cells; ICI, n=15 cells; DPDPE, n=17 cells; SNC80, n=14 cells; all across 3 biological replicates; dashed line indicates mean, shading +/− SEM). **(F)**. Nb39 max PM biosensor fluorescence significantly increases over baseline within 60 seconds of addition of either agonist DPDPE or SNC80 but not with addition of inverse agonist ICI, by one-way ANOVA (p<0.0001) with p-values from Dunnett’s multiple comparisons test to vehicle control reported in the figure. miniGsi max PM biosensor fluorescence significantly increases over baseline within 60 seconds of addition of either agonist DPDPE or SNC80 but not with addition of inverse agonist ICI, by one-way ANOVA (p<0.0001) with p-values from Dunnett’s multiple comparisons test to vehicle control reported in the figure. Venus-miniGs, a sensor for Gs coupling, fluorescence does not visibly increase after addition of 10µM SNC80. (Nb39: Ctrl, n=10 cells; ICI, n=17 cells; DPDPE, n=17 cells; SNC80, n=16 cells; miniGsi: Ctrl, n=17 cells; ICI, n=15 cells; DPDPE, n=17 cells; SNC80, n=14 cells; miniGs-SNC80, n=20 cells; all across 3 biological replicates; mean +/− 95% CI, points represent individual cells). **(G)**. Schematic of biosensor recruitment to DOR in intracellular compartments upon addition of a cell permeable agonist. Both Nb39 and miniGsi biosensors are diffuse throughout the cytoplasm in the absence of agonist (left), but are expected to localize to membranes containing active receptor upon agonist addition (right). **(H)**. PC12 cells expressing SNAP-DOR (magenta in merge) and Nb39-mVenus (green in merge) were treated with 10µM SNC80 and imaged live by confocal microscopy. Treatment with SNC80 leads to an increase in Nb39-mVenus signal in a perinuclear region (yellow arrow) which colocalizes with intracellular DOR (white in merge). A small amount of Nb39 recruitment is also visible at the PM (yellow arrowhead) (scale bar=5µm). **(I)**. PC12 cells expressing SNAP-DOR (magenta in merge) and Venus-miniGsi (green in merge) were treated with 10µM SNC80 and imaged live by confocal microscopy. miniGsi does not localize to intracellular DOR after agonist treatment, though a small amount of minGsi recruitment is visible at the PM (yellow arrowhead) (scale bar=5µm). **(J)**. Nb39 (solid line indicates mean, shading +/− SEM) and miniGsi (dashed line, shading +/− SEM) fluorescence in the region of the cell defined by intracellular DOR normalized to mean baseline fluorescence over time after addition of 10µM SNC80 (Nb39, n=49 cells across 4 biological replicates; miniGsi, n=51 cells across 3 biological replicates). **(K)**. Max intracellular biosensor fluorescence in the region of the cell defined by intracellular DOR within 120 seconds of agonist addition shows a significant increase in Nb39 recruitment with addition of permeable agonist SNC80 but not with peptide agonist DPDPE, by one-way ANOVA (p<0.0001) with p-values from Dunnett’s multiple comparison test to vehicle control reported in the figure. In contrast, miniGsi intracellular max fluorescence does not increase upon addition of 10µM SNC80 by one-tailed student’s t-test compared to vehicle control. miniGs intracellular max fluorescence also does not visibly increase upon SNC80 treatment. (Nb39: Ctrl, n=61 cells; DPDPE, n=61 cells; SNC80, n=49 cells; miniGsi: Ctrl, n=57 cells; SNC80, n=51 cells; miniGs: SNC80, n=36 cells; all across a minimum of 3 biological replicates; mean +/− 95% CI, points represent individual cells).

To test whether DOR localized to an intracellular compartment engages Nb39 or miniGsi differently upon activation by the same agonist, we took advantage of the fact that newly-synthesized DOR is retained in an intracellular compartment in neurons and PC12 cells (Fig. 1H, yellow arrow) acutely treated with nerve growth factor (NGF) (*16, 17*). We tested whether the two different biosensors were differentially recruited to DOR in intracellular compartments (IC DOR) vs. PM DOR. When cells were treated with 10µM SNC80, a membrane permeable, small molecule agonist, Nb39 was rapidly recruited to intracellular SNAP-tagged DOR, within 30 sec (Fig. 1H, Movie S1). When quantitated, Nb39 fluorescence in the region of the cell defined by intracellular DOR rapidly and significantly increased after SNC80 addition (Fig. 1J-K). This Nb39 recruitment was dynamic and required DOR activation, as the DOR antagonist Naltrindole rapidly reversed this effect (Fig. S2A-B). In striking contrast to Nb39, miniGsi was not recruited to IC DOR in the same time frame (Fig. 1I-J, Movie S2), despite comparable levels of IC DOR (Fig. S1B). MiniGsi fluorescence in the region of the cell defined by IC DOR did not increase in cells treated with SNC80 (Fig. 1J-K). Both sensors were also recruited to PM DOR, consistent with the TIR-FM results, although the sensitivity of detection was lower in confocal imaging (yellow arrowhead, Fig. 1H-I). As a control, miniGs was also not recruited to IC DOR in cells treated with SNC80 (Fig. 1K). The differential recruitment of two different active conformation biosensors to IC DOR suggests that agonist-induced activation of DOR in this intracellular compartment promotes an active receptor conformation recognized by only Nb39. Differential biosensor recruitment to IC DOR did not depend on the method used to cause DOR retention in this compartment (Fig. S3A), and recruitment to PM DOR was unaffected by the presence of IC DOR (Fig. S3B).

When cells were treated with 10µM peptide agonist DPDPE, which does not readily cross the PM over short time scales (*23*), Nb39 was not recruited to IC DOR (Fig. 1K). This suggests that activation of PM DOR is not sufficient for recruitment of Nb39 to IC DOR. We next tested whether PM DOR activation was required for differential sensor recruitment to IC DOR. Cells were pre-treated with a high concentration (100µM) of DOR inverse agonist ICI174864 (ICI), a peptide restricted to the extracellular space (*23*), to pharmacologically block PM DOR. Nb39 recruitment to IC DOR after 100nM SNC80 addition was then measured. Nb39 was robustly recruited to IC DOR, even when PM DOR was pharmacologically blocked, indicating that recruitment to IC DOR does not require activation of PM DOR (Fig 2A, top, 2B, C). When PM DOR was pharmacologically blocked, miniGsi again remained diffuse throughout the cell and was not recruited to IC DOR (Fig. 2A, bottom, 2B, C), indicating that the absence of miniGsi recruitment to IC DOR is not due to sequestration of sensor at PM DOR. To test whether activation of endogenous G proteins was restricting miniGsi recruitment, cells were pretreated with pertussis toxin (PTX) to inactivate endogenous Gαi/o proteins. Even in PTX-treated cells, miniGsi was not recruited to IC DOR, suggesting that competition with endogenous Gαi/o protein effectors for interaction with DOR is not responsible for the lack of recruitment of miniGsi to IC DOR (Fig. 2C).

**Fig. 2.**
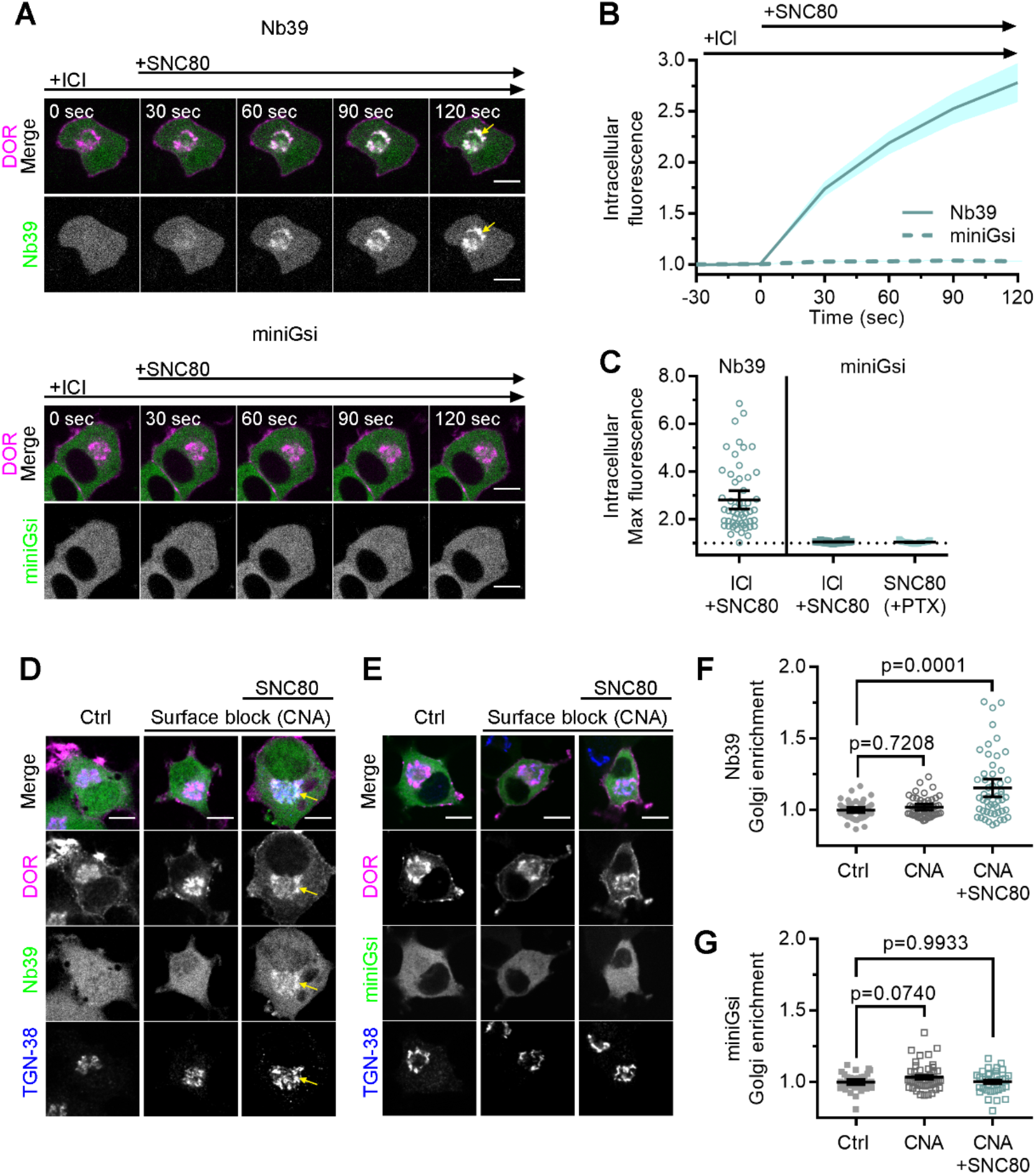
Differential sensor recruitment to Golgi DOR is independent of plasma membrane DOR activation. **(A)**. PC12 cells expressing SNAP-DOR (magenta in merge) and Nb39-mVenus or Venus-miniGsi (green in merge) were imaged live by confocal microscopy with 100µM ICI present in the media before addition of 100nM SNC80. After SNC80 treatment, Nb39-mVenus fluorescence increases in a perinuclear region (yellow arrow) which colocalizes with intracellular DOR (white in merge), whereas Venus-miniGsi remains diffuse through the cell (scale bar=5µm). **(B)**. Nb39 (solid line indicates mean, shading +/− SEM) or miniGsi (dashed line, shading +/− SEM) fluorescence in the region of the cell defined by intracellular DOR normalized to mean baseline fluorescence in cells treated with 100µM ICI and 100nM SNC80 (Nb39, n=54 cells across 3 biological replicates; miniGsi, n=58 cells across 4 biological replicates). **(C)**. Nb39-mVenus max intracellular fluorescence increases over baseline within 120 seconds of SNC80 in cells treated with 100µM ICI and 100nM SNC80. In contrast, miniGsi intracellular max fluorescence does not visibly increase over baseline in cells treated with ICI and SNC80, nor in cells pretreated with pertussis toxin (PTX) and SNC80 (Nb39: ICI+SNC80, n=54 cells; miniGsi: ICI+SNC80, n=58 cells; PTX+SNC80, n=33 cells; mean +/− 95% CI, points represent individual cells). **(D)**. PC12 cells expressing Flag-DOR and Nb39-mVenus or Venus-miniGsi, **(E)**, (green in merge) were treated with either 10µM β-chlornaltrexamine (CNA) alone for 15 minutes or 10µM CNA for 15 minutes followed by 10µM SNC80 for 5 minutes, then fixed and stained for Flag (magenta in merge) and *trans*-Golgi network marker TGN-38 (blue in merge) (scale bar = 5µm). Colocalization of DOR, Nb39, and TGN-38 is visible in white and light blue (yellow arrow) in cells treated with CNA and SNC80, but not CNA alone. **(F)**. Normalized Nb39-mVenus fluorescence enriched in the Golgi, expressed as sensor fluorescence in the region of the cell defined by the TGN-38 staining divided by sensor fluorescence in the region of the cell not containing TGN-38 staining. Nb39 Golgi enrichment is significantly increased in cells treated with CNA and SNC80, but not CNA alone, by one-way ANOVA (p<0.0001) with p-values reported in the figure from Dunnett’s multiple comparisons test compared to control cells (Ctrl, n=46 cells; CNA, n=49; CNA+SNC80, n=52; all across 2 biological replicates; points indicate individual cells with bars representing mean +/− 95% CI). **(G)**. Venus-miniGsi Golgi enrichment is not significantly increased in cells treated with either CNA and SNC80 or CNA alone, by one-way ANOVA (p=0.0654) with p-values reported in the figure from Dunnett’s multiple comparisons test compared to control cells (Ctrl, n=40 cells; CNA, n=50; CNA+SNC80, n=37; all across 2 biological replicates; points indicate individual cells with bars representing mean +/− 95% CI).

Immunofluorescence microscopy showed that Nb39 was recruited to IC DOR localized to the Golgi. Using a similar approach as described above, PC12 cells expressing Flag-tagged DOR and either Nb39 or miniGsi were pretreated for 15 minutes with 10µM β-chlornaltrexamine (CNA), an irreversible, cell impermeable antagonist (*17, 36*), to irreversibly block PM DOR, before treating with 10µM SNC80 for 5 minutes. Cells were stained for TGN-38, a marker for the trans-Golgi network, which was previously shown to colocalize with IC DOR (*17*). Consistent with live cell imaging data, only Nb39 and not miniGsi was recruited to IC DOR in a region of the cell colocalizing with the TGN-38 marker (Fig. 2D, E) in cells treated with CNA and SNC80. CNA alone did not cause recruitment of either sensor. Sensor fluorescence in the region of the cell defined by TGN-38 staining was normalized to sensor fluorescence in the cell outside this region, as a measure of sensor enrichment in the Golgi. Treatment with CNA and SNC80 significantly increased Nb39 Golgi enrichment (Fig. 2F), whereas miniGsi enrichment was not significantly different from control cells (Fig. 2G). These results confirm differential biosensor recruitment to IC DOR specifically localized to the Golgi and reiterate that PM DOR activation is not required for differential biosensor recruitment to Golgi DOR.

In addition to heterotrimeric G proteins, DOR and other GPCRs interact with other proteins and signaling effectors after agonist-induced conformational changes. Agonist-dependent differential biosensor recruitment to MOR and KOR in the PM correlates with recruitment of other receptor effectors, specifically G protein-coupled receptor kinase 2 (GRK2), which mediates receptor desensitization (*30*). Given differential recruitment of Nb39 and miniGsi to IC DOR, we hypothesized that Golgi localization may also influence coupling to other downstream signaling effectors. β-arrestins interact with DOR and other GPCRs after activation by agonists and receptor phosphorylation by GRKs to mediate receptor desensitization and internalization from the PM (*37*–*39*). β-arrestins can also scaffold kinase signaling complexes from GPCRs at the PM and endosomes (*40*–*44*).

To test whether β-arrestin effectors are recruited to active IC DOR, we monitored recruitment of fluorescently tagged β-arrestin1 or β-arrestin2 to IC DOR. Similar to miniGsi, neither β-arrestin-1 nor β-arrestin-2 were recruited to IC DOR in cells treated with SNC80, and no increase in fluorescence in the region of the cell defined by IC DOR was detected (Fig. 3A-C). In contrast, both β-arrestins were visibly recruited to the PM by confocal imaging (Fig. 3A, B, yellow arrowheads) and by quantitation of TIR-FM imaging (Fig. 3D-F) in response to SNC80 treatment. Together, these data indicate that like miniGsi, β-arrestins interact with only agonist-activated DOR present in the PM.

**Fig. 3.**
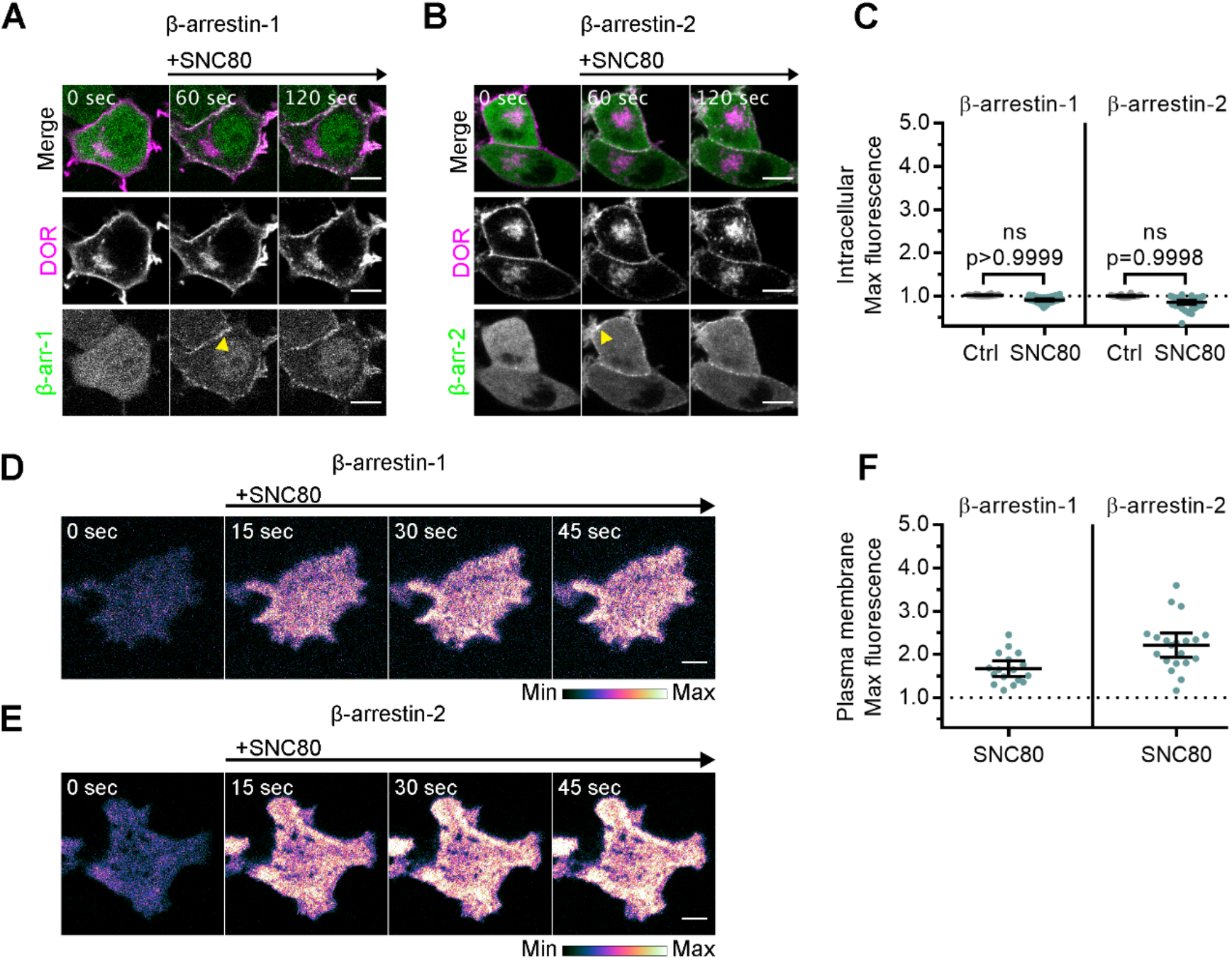
Arrestins are differentially recruited to plasma membrane and intracellular DOR. **(A)**. PC12 cells expressing SNAP-DOR (magenta in merge) and β-arrestin-1-mScarlet (green in merge) were treated with 10µM SNC80 and imaged live by confocal microscopy. β-arrestin-1-mScarlet signal increases at the PM (yellow arrowhead) but not at sites colocalized with intracellular DOR upon 10µM SNC80 treatment (scale bar=5µm). **(B)**. PC12 cells expressing SNAP-DOR (magenta in merge) and β-arrestin-2-tdTomato (green in merge). β-arrestin-2-tdTomato signal increases at the PM (yellow arrowhead) but not at sites colocalized with intracellular DOR upon 10µM SNC80 treatment (scale bar=5µm). **(C)**. Neither β-arrestin-1-mScarlet nor β-arrestin-2-tdTomato max intracellular fluorescence significantly increases within 120 seconds of SNC80 addition by one-tailed student’s t-test compared to control cells (β-arr-1: Ctrl, n=16 cells; SNC80, n=33 cells; β-arr-2: Ctrl, n=14 cells; SNC80, n=37 cells; with control conditions across 1 biological replicate and SNC80 conditions across 3 biological replicates; mean +/− 95% CI, points represent individual cells). **(D)**. β-arrestin-1-mScarlet in PC12 cells expressing SNAP-DOR imaged using TIR-FM to capture recruitment to the PM after addition of 10µM SNC80 (scale bar = 5µm). **(E)**. β-arrestin-2-tdTomato in PC12 cells expressing SNAP-DOR imaged using TIR-FM to capture recruitment to the PM after addition of 10µM SNC80 (scale bar = 5µm). Calibration bars indicate relative fluorescence values in scaled images. **(F)**. Both β-arrestin-1-mScarlet and β-arrestin-2-tdTomato max PM fluorescence increases within 60 seconds of 10µM SNC80 addition (β-arr-1: SNC80, n=17 cells; β-arr-2: SNC80, n=20 cells; all across 3 biological replicates; mean +/− 95% CI, points represent individual cells).

Given differential recruitment of active conformation biosensors, Nb39 and miniGsi, and the absence of β-arrestin recruitment to IC DOR, we asked whether the active conformation of IC DOR allows for signaling through G proteins. Like the other opioid receptors, DOR couples primarily to Gαi/o proteins which inhibit adenylyl cyclase activity to decrease cAMP (*45*). We used a Forster resonance energy transfer (FRET) sensor, ICUE3, to monitor cAMP levels in single cells in real time (*46*). In PC12 cells expressing ICUE3 and SNAP-DOR, addition of adenylyl cyclase activator forskolin (Fsk, 2µM) caused a rapid increase in the CFP/FRET ratio over the baseline ratio (Fig, 4A-B). Pretreatment with 100nM SNC80 prior to Fsk addition decreased the Fsk-stimulated cAMP response (Fig. 4C-D), consistent with DOR activation inhibiting adenylyl cyclase activity.

**Fig. 4.**
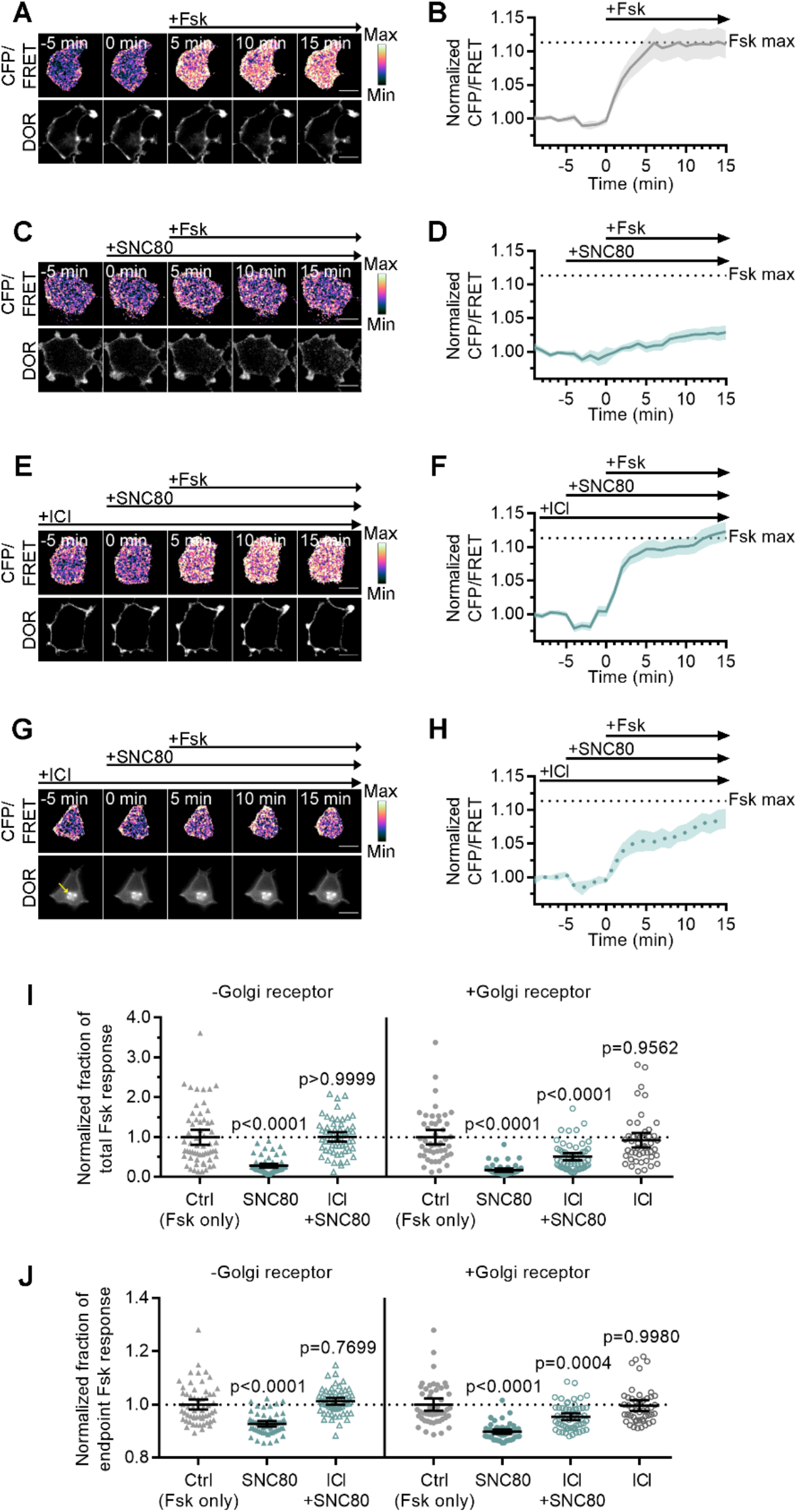
Golgi DOR inhibits cAMP. **(A-H)**. Ratiometric CFP/FRET and receptor images, along with corresponding trace of mean cellular CFP/FRET ratios (solid line indicates mean, shading +/− 95% CI), in PC12 cells expressing the ICUE3 cAMP FRET sensor and SNAP-DOR (scale bar=10µm). Calibration bars indicate relative fluorescence values in scaled images. **(A-B)**. In cells without intracellular DOR, CFP/FRET ratio increases over baseline upon treatment with 2µM forskolin (Fsk), consistent with increase in cellular cAMP levels. **(C-D)**. Treatment with DOR agonist SNC80 (100nM) decreases Fsk-stimulated increase in CFP/FRET ratio. **(E-F)**. In cells without intracellular DOR, SNC80-dependent decrease in Fsk-stimulated CFP/FRET ratio is reversed when peptide inverse agonist ICI (100µM) is present in media. **(G-H)**. In cells containing intracellular DOR (**G**, yellow arrow), SNC80 decreases Fsk-stimulated CFP/FRET ratio even when ICI is present in media. **(I-J)**. Fsk-stimulated total cAMP levels (area under the curve) **(I)** and endpoint CFP/FRET ratios **(J)**, normalized to mean of control treated cells within -Golgi receptor and +Golgi receptor groups. Treatment with 100nM SNC80 significantly decreases total Fsk-stimulated cAMP and endpoint ratios. ICI and SNC80 treatment of cells without Golgi DOR does not significantly decrease total cAMP or endpoint ratios. In contrast, ICI and SNC80 treatment of cells with Golgi DOR significantly decreases total cAMP and endpoint ratios. ICI treatment alone in cells with Golgi DOR does not significantly decrease total cAMP and endpoint ratios. (-Golgi receptor: control, n= 59 cells; SNC80, n=58; ICI+SNC80, n=57; +Golgi DOR: control, n=48; SNC80, n=50; ICI+SNC80, n=55; ICI, n=47; all across 2 biological replicates; one-way ANOVA (total cAMP, p<0.0001; endpoint cAMP, p<0.0001) with p-values reported in the figure from Sidak’s multiple comparisons test for each condition compared to control cells within –Golgi receptor and +Golgi receptor groups).

To specifically test whether IC DOR in the Golgi is sufficient for cAMP inhibition, we blocked PM DOR with the peptide inverse agonist ICI in cells either acutely treated or untreated with NGF to induce retention of DOR in the Golgi. The overall Fsk response and SNC80-mediated inhibition in cells treated with NGF were comparable to untreated cells (Fig. S4A). To isolate the contribution of IC DOR to cAMP inhibition, we pharmacologically blocked PM DOR with 100µM ICI. In cells with PM DOR only, SNC80 failed to decrease Fsk-stimulated cAMP in the presence of ICI (Fig. 4E-F). Endpoint and total cAMP levels were not significantly different from cells treated with Fsk alone (Fig. 4I-J). In contrast, in cells with IC DOR, SNC80 decreased Fsk-stimulated cAMP even in the presence of ICI (Fig. 4G-H), and significantly decreased endpoint and total cAMP levels (Fig. 4I-J). As a control, ICI alone did not significantly affect endpoint or total cAMP levels (Fig. 4I-J). These results indicate that Golgi DOR activation is sufficient for cAMP inhibition.

As an independent method to induce an intracellular pool of DOR, we used LY294002, a small molecule inhibitor of PI3K that causes DOR retention in the Golgi independent of NGF (*47*). Similar to results obtained in NGF-treated cells, the permeable small molecule agonist SNC80 decreased Fsk-stimulated cAMP (Fig. S4B-D) even in the presence of ICI, reiterating that Golgi DOR activation is sufficient for cAMP inhibition. The endpoint and total cAMP levels decreased significantly compared to cells treated with Fsk alone (Fig. S4G-H). As a control, the peptide DPDPE agonist did not decrease the Fsk response in the presence of ICI (Fig. S4B, E-F).

## Discussion

Together, our data suggest that DOR activation by the *same* agonist in *different* subcellular compartments promotes distinct active conformations recognized differentially by biosensors. The possibility that Nb39 and miniGsi could recognize distinct conformations is supported by structures of agonist-bound homologous MOR and KOR in complex with Nb39 or MOR in complex with the heterotrimeric G protein complex, Gα_i1_β_1_γ_2_. Structures of MOR with agonist BU72 and KOR with agonist MP1104 share the outward shift of transmembrane helix (TM) 6 which is characteristic of active GPCR structures (*32, 33*). Nb39 appears to stabilize this conformation via contacts with intracellular loops (ICL) 2 and ICL3, as well as the 8^th^ helix through residues conserved across MOR, KOR, and DOR (*32, 33*). The structure of MOR in complex with agonist DAMGO and the nucleotide free Gαi protein is very similar to the MOR-Nb39 structure with the exception of a greater displacement of TM6 toward TM7 and decreased extension of ICL3 (*48*). The miniGsi sensor does not contain all regions of Gαi which contact the receptor, but many of the residues which interact with MOR ICL2 and ICL3 via the Gαi C-terminal α5 helix are present in miniGsi, and previous reports show that miniGs and Gαs contact B2AR similarly (*34, 49*). Though Nb39 and Gαi contact opioid receptors in similar regions and share two interaction residues, each also makes additional distinct contacts with TM domains and the 8^th^ helix. Distinct interactions with these intracellular domains important for effector coupling and unique stabilization of TM6 and ICL3 by Nb39 could suggest that these sensors differentially report distinct conformations relevant to receptor function. Additionally, these structures are limited to a single static view of opioid receptor active conformation, and the ability of Nb39 and miniGsi to discriminate between additional distinct intermediate or active conformations will be an exciting area for future study.

One clear difference between compartments is the composition of specific phospholipids that make up the membranes (*50*–*52*). Phospholipids differing in charge can stabilize active or inactive conformations of the β_2_-adrenergic receptor, stabilize G protein coupling, and modulate G protein selectivity (*53*–*55*). Lipid composition can also influence recruitment of effectors like β-arrestins which bind PI4,5P2, a phospholipid species enriched in the plasma membrane (*56*), potentially contributing to the lack of observed arrestin recruitment to Golgi DOR. To date, β-arrestin recruitment to active GPCRs in the Golgi has not been reported, and the impact of receptor localization to this compartment on desensitization, β-arrestin recruitment, and β-arrestin biased signaling is not known.

Other compartment-specific factors including ion concentrations and GPCR interacting proteins could influence receptor conformations and effector coupling. The Golgi lumen is both more acidic than the extracellular space, pH 6.4 vs pH 7.4 (*57*–*59*), which could affect ligand binding and GPCR activation (*60*–*63*). Estimated concentrations of sodium in the Golgi are closer to cytosolic sodium concentrations (12-27mM) than high extracellular sodium concentrations (100mM) (*64, 65*). Sodium acts as an allosteric modulator of class A GPCRs, and DOR specifically has been crystallized with a coordinated sodium ion, which stabilizes the inactive receptor conformation and is required for receptor activation and signaling (*66, 67*), suggesting Golgi sodium concentrations could affect DOR activity. Additionally, DOR interacting proteins, especially those which regulate DOR trafficking and localize to the Golgi, like the COPI complex and Rab10, could also regulate receptor conformations and effector coupling (*68*–*70*).

Our results suggest that the conformational space sampled by any given GPCR, even when activated by the same agonist, differs based on the precise subcellular location of the receptor. Compartmental effects on GPCR conformations may also be specific to individual GPCRs. In contrast to DOR, both an active state nanobody and miniGs are recruited to active Gαs-coupled β_1_-adrenergic receptor (B1AR) in the Golgi, suggesting the local Golgi environment may influence DOR and B1AR energy landscapes differently (*71, 72*). These GPCR specific effects may reflect important differences in pharmacology among individual GPCRs and emphasize the importance of characterizing compartmental effects for each GPCR.

These results also provide a new perspective into drug development efforts, by highlighting the effects that the subcellular location of receptors could have on the integrated effects of any given drug. The majority of these efforts largely rely on assays using conventional readouts of signaling in model cells, where GPCR localization could be different from that of physiologically relevant cells *in vivo*. This difference is especially true for DOR, which exhibits robust surface localization in model cell lines, but high levels of intracellular pools in many neuronal subtypes. Traditional signaling assays, which rely on whole-cell readouts of primary signaling pathways such as cAMP, will not distinguish between the contributions of different pools of receptors, which could signal differently via pathways outside the primary readouts (*73*). Therefore, the potentially distinct effects of ligands at spatially distinct pools of receptors in the integrated response should be an important consideration for measuring the outcomes of receptor activation.

## Materials and Methods

### DNA constructs

SSF-DOR construct consists of an N-terminal signal sequence followed by a Flag tag followed by the mouse DOR sequence in a pcDNA3.1 vector backbone. To create SNAP-DOR, the full-length receptor sequence was amplified from the SSF-DOR construct by PCR with compatible cut sites (BamHI and XbaI). The SNAP tag (New England Biolabs, Ipswich, MA) was amplified by PCR with compatible cut sites (HindIII and BamHI) and both were ligated into a pcDNA3.1 vector backbone to produce the final construct containing an N-terminal signal sequence, followed by the SNAP tag and then the receptor. β-arrestin-1 was generated from a geneblock (Integrated DNA Technologies, Coralville, IA) containing the human cDNA (ENST00000420843) for hARRB1 with HindIII and AgeI cut sites. mScarlet was amplified by PCR from pmScarlet_alphaTubulin_C1 a gift from Dorus Gadella (Addgene plasmid #85045) (*74*), with AgeI and XbaI cut sites. Both were then ligated into a pcDNA3.1 vector backbone to produce a C-terminally tagged β-arrestin-1. β-arrestin 2 tagged with tdTomato was generated from β-arrestin 2-GFP via restriction site cloning (*44*). Nb39-mVenus was a gift from Drs. Bryan Roth and Tao Che. Venus-miniGsi and Venus-miniGs were gifts from Drs. Greg Tall and Nevin Lambert. pcDNA3-ICUE3 was a gift from Dr. Jin Zhang (Addgene plasmid #61622) (*46*).

### Cell culture and transfection

Pheochromocytoma-12 cells (PC12 cells, ATCC #CRL-1721) were used for all experiments. Cells were maintained at 37°C with 5% CO2 and culture in F-12K media (Gibco, #21127), with 10% horse serum and 5% fetal bovine serum (FBS). Cells were grown in flasks coated with CollagenIV (Sigma-Aldrich, #C5533) to allow for adherence. PC12 cells were transiently transfected at 90% confluency according to manufacturer’s guidelines with Lipofectamine 2000 (Invitrogen, #11668) with 1.5ug of each DNA construct to be expressed. The transfection mixture was incubated with cells in Opti-MEM media (Gibco, #31985) for 5 hours then removed and replaced with normal culture media until imaging 48-72 hours following transfection.

### Live cell imaging with fluorescent biosensors

PC12 cells transfected with SNAP-DOR and the appropriate biosensor were plated and imaged in single-use MatTek dishes (MatTek Life Sciences, #P35G-1.5-14-C) coated with 20μg/mL poly-D-lysine (Sigma-Aldrich, #P7280) for 1 hr. For experiments requiring a Golgi pool of DOR, cells were pretreated with 100ng/mL of NGF (Gibco, #13257) or 10μM LY294002 (Tocris, #1130) or 20μM PI4K inhibitor MI 14 (Tocris, #5604) for 1 hour prior to imaging, as described previously (*16, 17, 22*). Cells were labeled with 500nM SNAP-Surface 649 (New England Biolabs, #S9159S) for 5 minutes at 37°C for TIR-FM imaging or 1μM permeable SNAP-Cell 647-SiR (New England Biolabs, #S9102S) for 15 minutes followed by a 15 minute wash in cell culture media for confocal imaging. Cells were imaged on a Nikon TiE inverted microscope using a 60x/1.49 Apo-TIRF (Nikon Instruments, Melville, NY) objective in CO_2_-independent Leibovitz’s L-15 media (Gibco, #11415), supplemented with 1% FBS in a 37°C heated imaging chamber (In Vivo Scientific). RFP (β-arrestin-1 and β-arrestin-2, 561 nm excitation, 620 emission filter), YFP (Nb39-mVenus and Venus-miniGsi, 488nm excitation, 446/523/600/677 quad-band filter) and the SNAP labeled DOR (647nm excitation, 700 emission filter when imaged with RFP or 446/523/600/677 quad-band filter when imaged with YFP) were excited with solid state lasers and collected with an iXon + 897 EMCCD camera (Andor, Belfast, UK).

### Immunofluorescence and fixed cell imaging

PC12 cells transfected with Flag-DOR and either Nb39-mVenus or Venus-miniGsi were plated on poly-D-lysine coated coverslips and grown at 37°C for 48 hours. To induce intracellular accumulation of newly synthesized DOR, cells were treated with NGF (100ng/mL) for 1 hour prior to treatment for 15 minutes with 10µM β-chlornaltrexamine (CNA, Sigma-Aldrich, #O001) or CNA followed by 10µM SNC80 (Tocris, #0764) for 5 minutes. Cells were fixed with 4% paraformaldehyde, pH 7.4, for 20 minutes at 25°C followed by blocking with phosphate-buffered saline (PBS) with 5% fetal bovine serum (FBS), 5% glycine, 0.75% Triton-X-100, 1mM magnesium chloride, and 1mM calcium chloride. Primary and secondary antibody incubations were performed for 1 hour at 25°C in blocking buffer with anti-Flag-M1 (Sigma-Aldrich, #F3040, 1:1000) conjugated with Alexa-647 (Molecular Probes, #A20186) and anti-TGN-38 rabbit polyclonal antibody (Sigma-Aldrich, #T9826, 1:1000), and goat anti-Rabbit IgG conjugated to Alexa-568 (ThermoFisher, #A-11011, 1:1000), respectively. Cells were washed with blocking buffer without Triton-X-100 after primary and secondary incubations. Coverslips were mounted on glass slides using Prolong Diamond Reagent (Molecular Probes, #P36962). Cells were imaged on a Nikon TiE inverted microscope using a 60x/1.49 Apo-TIRF (Nikon Instruments, Melville, NY) objective and iXon + 897 EMCCD camera (Andor, Belfast, UK).

### Live cell FRET imaging

PC12 cells transfected with SNAP-DOR and the ICUE3 FRET sensor were plated and imaged in MatTek dishes (MatTek Life Sciences, #P35G-1.5-14-C) coated with poly-D-lysine. For experiments comparing inhibition of Fsk-stimulated cAMP in cells with and without Golgi DOR, cells in all conditions were treated with cycloheximide (3μg/mL) for 1 hour before imaging to chase out any receptor transiting through the biosynthetic pathway. To induce a Golgi pool, cells were treated with NGF for 1 hour prior to cycloheximide treatment to build up a pool of internal receptors which is maintained even in the absence of new protein synthesis with NGF maintained in the media during the subsequent cycloheximide incubation (*17*). Cycloheximide, NGF, and 100μM ICI174864 (ICI, Tocris, #0820), when appropriate, were present in the media for the duration of the experiment. For experiments comparing the signaling of DPDPE (Tocris, #1431) peptide agonist and SNC80 small molecule agonist with and without a surface block, cells in all conditions were treated with 10μM LY294002 for 1hr prior to imaging to induce a Golgi pool of receptor, and LY294002 was maintained in the media throughout the duration of the experiment. Cells were labeled with 1μM permeable SNAP-Cell 647-SiR for 15 minutes followed by a 15 minute wash to visualize receptor. Cells were imaged in L-15 media supplemented with 1% FBS at 25°C in a temperature-controlled imaging chamber (In Vivo Scientific) at 60 second intervals. Imaging was conducted on a Ti2 inverted microscope (Nikon Instruments, Melville, NY) with a 60x NA 1.49 Apo-TIRF objective (Nikon Instruments, Melville, NY). CFP (405 nm excitation, 400 emission filter), YFP or FRET (405nm excitation, 514 emission filter) and the SNAP tagged isoform (647-nm excitation, 700 emission filter) were collected with a iXon-888 Life EMCCD camera (Andor, Belfast, UK) every 30 sec with 5 frames of baseline before 2μM Fsk (Sigma-Aldrich, #F3917) addition to stimulate adenylyl cyclase activity.

### Image quantification

All image quantification was performed using ImageJ/Fiji (National Institutes of Health, Bethesda, MD) (*75, 76*). To quantify biosensor recruitment to intracellular DOR or plasma membrane DOR, the receptor channel at each timepoint was thresholded and used to create a binary mask to isolate only pixels containing receptor signal. The receptor mask from each timepoint was then applied to the corresponding timepoint in the biosensor channel to produce an image of biosensor fluorescence in regions of the cell containing receptor. A region of interest corresponding to intracellular receptor was selected in confocal images, and in TIRF images a region of interest capturing the entire cell was selected. Mean fluorescence intensity was then measured in these images over time and normalized to average baseline fluorescence before drug addition.

A similar approach was used to measure biosensor recruitment to the Golgi in cells fixed and stained for TGN-38. In these images, TGN-38 was used to create the binary mask, which was then applied to the biosensor channel to isolate biosensor fluorescence in the Golgi region of the cell. An inverse mask of the TGN-38 channel was also created and applied to the biosensor channel to isolate biosensor fluorescence in all other regions of the cell. Biosensor enrichment in the Golgi is expressed as the mean fluorescence intensity in the Golgi region, divided by the mean fluorescence intensity in the rest of the cell. FRET images were analyzed in ImageJ as previously described (*17, 44*). Briefly, the CFP channel was divided by the FRET channel at each timepoint. A region of interest was defined for each cell in a given field and the resulting CFP/FRET ratio measured at each timepoint. The CFP/FRET ratio was normalized to the mean baseline ratio before drug addition for each cell. Endpoint CFP/FRET ratios and total cAMP responses (area under the curve) for all cells were normalized to the average of cells in the control condition for each experimental replicate.

## Acknowledgements

The authors would like to thank Dr. Alan Smrcka, Dr. Lois Weisman, Dr. Bing Ye, and Candilianne Serrano Zayas for valuable feedback on this project. We also thank Drs. Bryan Roth, Tao Che, Greg Tall, and Nevin Lambert for essential reagents.

## Author Contributions

SEC and MAP conceptualized the research project. SEC, WK, ZYW, and MAP participated in research design. SEC and WK performed experiments and collected and analyzed data. SEC, WK, and ZYW contributed new reagents or analytic tools. SEC, WK, ZYW, and MAP contributed to writing of the paper. MAP supervised the research.

