## Supplemental figures and legends for "Conformational specificity of opioid receptors is determined by subcellular location irrespective of agonist"

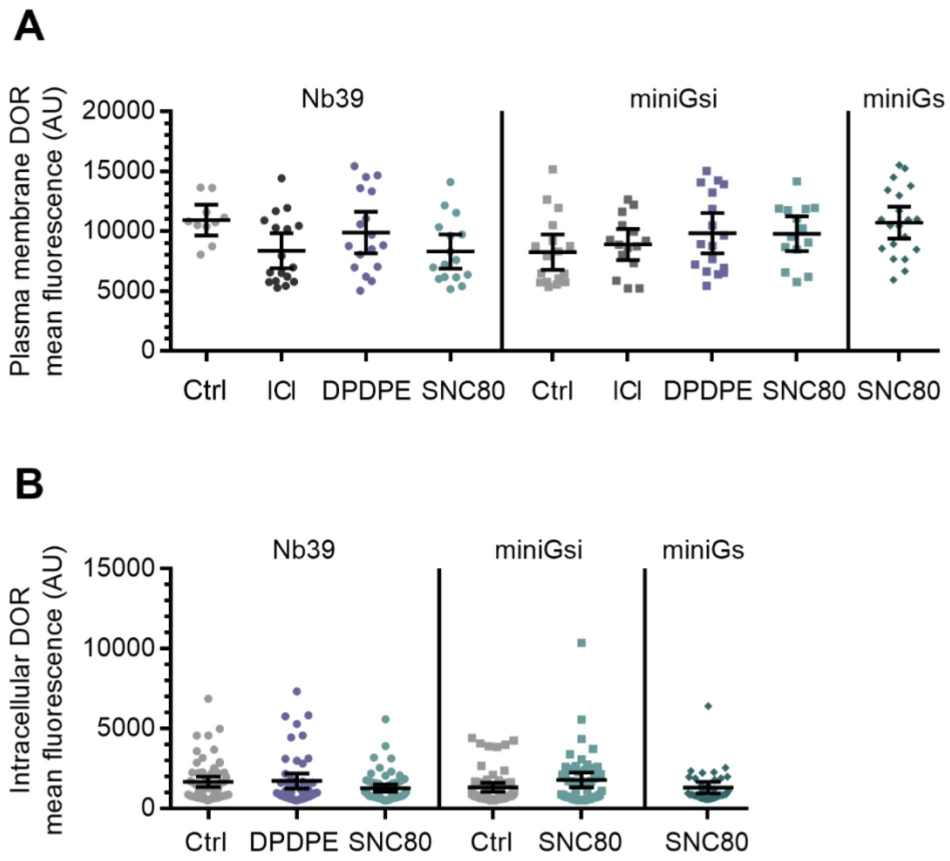


**Fig. S1. DOR expression levels are similar across treatment conditions.**

**(A).** Plasma membrane DOR expression levels in cells expressing Nb39, miniGsi, or miniGs are similar across untreated or 10µM agonist treatment conditions control (Nb39: Ctrl, n=10 cells; ICI, n=17 cells; DPDPE, n=17 cells; SNC80, n=16 cells; miniGsi: Ctrl, n=17 cells; ICI, n=15 cells; DPDPE, n=17 cells; SNC80, n=14 cells; miniGs-SNC80, n=20 cells; across a minimum of 3 biological replicates; mean +/- 95% CI, points represent individual cells). **(B).** Intracellular DOR expression levels in cells expressing Nb39, miniGsi, or miniGs are similar across untreated or 10µM agonist treatment conditions (Nb39: Ctrl, n=61 cells; DPDPE, n=61 cells; SNC80, n=49 cells; miniGsi: Ctrl, n=57 cells; SNC80, n=51 cells; miniGs: SNC80, n=36 cells; all across 3 biological replicates; mean +/- 95% CI, points represent individual cells).

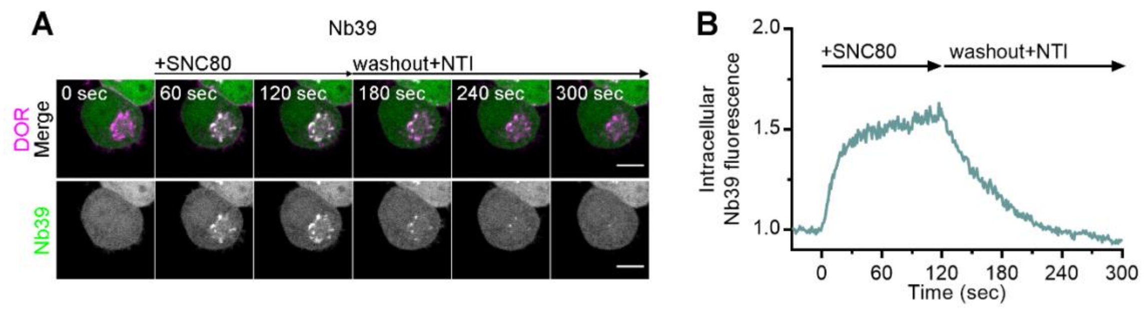


**Fig. S2. Nb39 recruitment to active DOR is reversible.**

**(A).** PC12 cells expressing SNAP-DOR (magenta in merge) and Nb39-mVenus (green in merge) were imaged live by confocal microscopy. Cells were pretreated with irreversible, impermeable antagonist CNA (1µM) for 15 minutes prior to imaging to inhibit plasma membrane DOR activation and internalization. After 1µM SNC80 treatment, Nb39-mVenus fluorescence increases in a perinuclear region which colocalizes with intracellular DOR (white in merge), and this recruitment is reversed upon a washout, introducing fresh imaging media containing permeable antagonist naltrindole (NTI, 10µM) (scale bar=5µm). **(B).** Representative trace of Nb39 fluorescence in the region of the cell defined by intracellular DOR normalized to mean baseline fluorescence for the cell shown in **(A).**

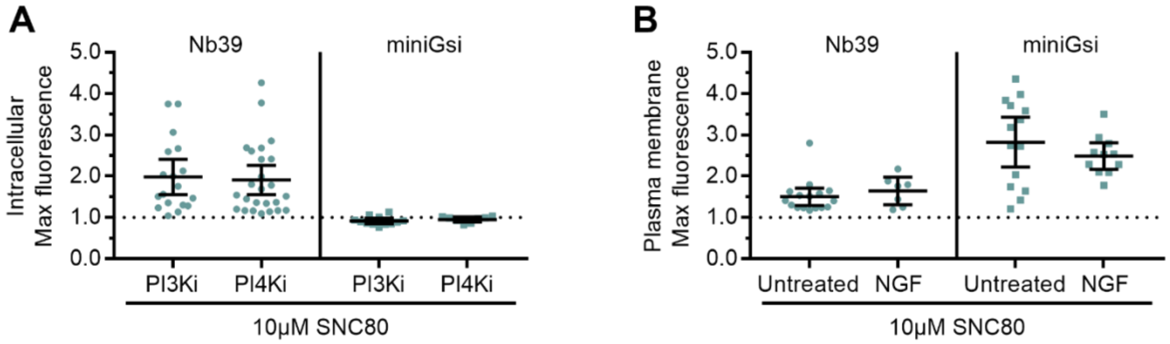


**Fig. S3. Mechanism of DOR Golgi retention does not influence sensor recruitment to Golgi or PM DOR.**

**(A).** Nb39, but not miniGsi, max intracellular fluorescence increases over baseline within 120 seconds of 10μM SNC80 addition in cells treated with 10μM PI3K inhibitor LY294002 or 20μM PI4K inhibitor MI14 to induce DOR retention in the Golgi through a DOR specific and non-specific mechanism (*77, 78*), respectively (Nb39: PI3Ki, n=18 cells; PI4Ki, n=25 cells; miniGsi: PI3Ki, n=12 cells; PI4Ki, n=8 cells; all across 1 biological replicate; mean +/- 95% CI, points represent individual cells). **(B).** Nb39 and miniGsi max plasma membrane fluorescence increases to a similar degree within 60 seconds of 10μM SNC80 addition in untreated cells without Golgi DOR and NGF-treated cells with Golgi DOR (Nb39: Untreated, n=17 cells;  NGF, n=7 cells; miniGsi: Untreated, n=14cells; NGF, n=11 cells; all across 1 biological replicate; mean +/- 95% CI, points represent individual cells).

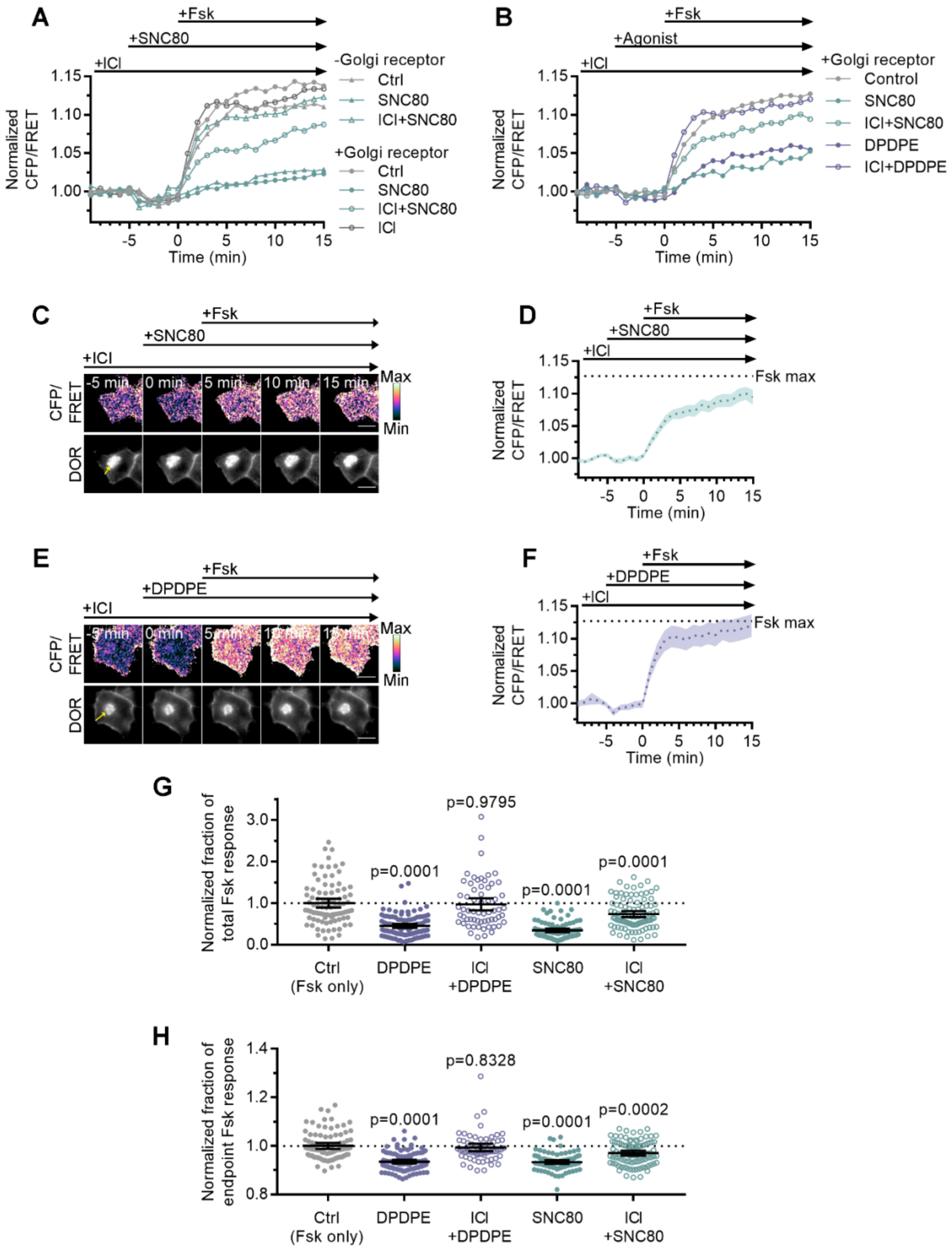


**Fig. S4. Golgi DOR inhibits cellular cAMP.**

**(A).** Trace of mean cellular cAMP levels in PC12 cells expressing SNAP-DOR and cAMP FRET sensor ICUE3 with and without Golgi receptor. (-Golgi receptor: control, n= 59 cells; SNC80, n=58; ICI+SNC80, n=57; +Golgi DOR: control, n=48; SNC80, n=50; ICI+SNC80, n=55; ICI, n=47). **(B).** Trace of mean cellular cAMP levels in PC12 cells expressing SNAP-DOR and ICUE3 with Golgi receptor and treated with either small molecule agonist SNC80 or peptide agonist DPDPE. (control, n=89 cells; DPDPE, n=101; ICI+DPDPE, n=62; SNC80, n=87; ICI+SNC80, n=95). **(C-F).** Ratiometric CFP/FRET and receptor images, along with corresponding trace of mean cellular CFP/FRET ratios (solid line indicates mean, shading +/- 95% CI), in PC12 cells expressing the ICUE3 cAMP FRET sensor and SNAP-DOR (scale bar=10µm). Calibration bars indicate relative fluorescence values in scaled images. **(C-D).** In cells containing Golgi DOR (**E**, yellow arrow), SNC80 (100nM) decreases Fsk-stimulated cAMP levels even when peptide inverse agonist ICI (100µM) is present in media. **(E-F).** In cells containing Golgi DOR (**C**, yellow arrow), peptide agonist DPDPE (100nM) does not decrease Fsk-stimulated cAMP levels when peptide inverse agonist ICI (100µM) is present in media. **(G-H).** Fsk-stimulated total cAMP levels (area under the curve) **(G)** and endpoint CFP/FRET ratios **(H),** normalized to the mean of control treated cells. Cells in all conditions have intracellular DOR. Total and endpoint cAMP responses are significantly decreased in cells treated with 100nM peptide agonist DPDPE or small molecule agonist SNC80. Total and endpoint cAMP responses are significantly decreased only in cells treated with membrane permeable agonist SNC80 and not DPDPE when 100µM ICI is present in media. (Ctrl, n=89 cells; DPDPE, n=101; ICI+DPDPE, n=62; SNC80, n=87; ICI+SNC80, n=95; all across 3 biological replicates; one-way ANOVA (total, $p<0.0001$; endpoint, $p<0.0001$) with p-values reported in the figure from Dunnett's multiple comparison test for each condition compared to control (Fsk only) condition).

**Movie S1.**

In PC12 cells expressing SNAP-DOR (magenta), Nb39 (green) is rapidly recruited to intracellular DOR (white) upon addition of 10µM SNC80 (Scale bar=5µm).

**Movie S2.**

In PC12 cells expressing SNAP-DOR (magenta), miniGsi (green) is not recruited to intracellular DOR upon addition of 10µM SNC80 (Scale bar=5µm).
